# Human landscape modification shapes foraging preferences and sucrose responsiveness of honey bees in Asia

**DOI:** 10.64898/2026.02.27.708048

**Authors:** Gifty Alin Jacob, Manish Ravi, Sachin Bhaskar, Abhinay Arra, Hema Somanathan, Ingolf Steffan-Dewenter, Ricarda Scheiner

**Affiliations:** Behavioral Physiology and Sociobiology, Biocenter, Julius-Maximilians-Universität Würzburg, Würzburg, Germany; School of Biology, Indian Institute of Science Education and Research Thiruvananthapuram, Thiruvananthapuram, Kerala, India; Animal Ecology and Tropical Biology, Biocenter, Julius-Maximilians-Universität Würzburg, Würzburg, Germany

**Keywords:** *Apis*, Asian honey bees, Division of foraging labour, Nectar preferences, Resource quality

## Abstract

Landscape composition is central in shaping how pollinators utilise floral resources, yet its influence on foraging behaviour of co-occurring Asian honey bees remains underexplored. Resolving this gap is crucial to understand how closely-related, native and introduced species maintain foraging efficiency in rapidly changing environments. We investigated nectar preferences, sucrose responsiveness, and foraging task partitioning in three co-occurring honey bee species in India: *Apis florea* (native open-nesting), *Apis cerana* (native cavity-nesting), and *Apis mellifera* (introduced cavity-nesting), across forest, agricultural, and urban landscapes. Landscape type strongly influenced crop sugar concentrations of honey bees. While all species collected high-concentration nectar in forests, *A. mellifera* and *A. cerana* collected lower concentrations than *A. florea* in urban habitats. *A. florea* showed consistent preference for high-concentration nectar across landscapes. Complementing this, sucrose responsiveness assays revealed a lower responsiveness of *A. florea* compared to cavity-nesting species. Foraging task partitioning differed among species, but interestingly, also among landscapes. While *A. cerana* predominantly collected nectar, *A. mellifera* foraged equally for pollen, nectar and water, and *A. florea* shifted task allocation across landscapes. In conclusion, we provide the first comparative evidence that landscape composition and species characteristics jointly shape foraging preferences and organisation of foraging labour in Asian honey bees.

## INTRODUCTION

Pollinators play an essential role in maintaining ecosystem functioning and ensuring global food security [1]. However, their foraging success is increasingly challenged by rapid land-use change and habitat degradation. Landscape composition fundamentally shapes the foraging ecology of pollinators by determining the abundance, diversity, and accessibility of floral resources [2–7]. Changes in the habitat structure and land use can impact the quality of resources, such as nectar sugar concentration and pollen protein content, and thus the profitability of foraging sites [8,9]. Shifts in the landscape context can significantly affect key pollinators such as honey bees, with direct consequences on their colony health and survival, as reductions in high-quality resources can constrain colony nutrition and fitness [10,11]. Evidence from both temperate and tropical systems shows that foraging distances, activity, recruitment behaviour, and even colony growth in honey bees are closely linked to landscape composition and habitat diversity [5,12–15]. Moreover, different social bee taxa often respond idiosyncratically to landscape gradients, as seen in the contrasting responses of the Eastern honey bee, *Apis cerana*, and the stingless bee *Tetragonula laeviceps* to land-use change [8], underscoring the interplay between environmental context and species-specific ecology.

Foraging is a central decision-making process in animals, shaped by the need to maximise nutrient intake while minimising energetic costs [16,17]. In social insects, these individual decisions are integrated across thousands of workers, collectively determining colony nutrition, reproduction, and survival [18,19]. Eusocial colonies achieve such efficiency through division of labour, where specialised workers perform complementary tasks [20–22]. In honey bees, this includes specialisation among foragers collecting nectar, pollen, or water (table 1) [23–26]. Task allocation is closely linked to sensory responsiveness, particularly sucrose responsiveness, which influences an individual’s likelihood of foraging for specific resources [25–29]. Workers with high sucrose responsiveness (i.e. low response thresholds for sugar water) tend to collect pollen or water, while those with lower responsiveness forage for nectar [25,28]. This relationship makes sucrose responsiveness a reliable behavioural indicator of context-dependent foraging preference and task differentiation [29–31].

**Table 1:** Foraging roles classified based on the characteristics of the material collected and the crop content of foragers [23–25].

| Foraging role | Pollen | Crop volume > 1 µl | Crop concentration > 5% Brix |
| --- | --- | --- | --- |
| Pollen | ✓ | ✗ | ✗ |
| Pollen and water | ✓ | ✓ | ✗ |
| Pollen and nectar | ✓ | ✓ | ✓ |
| Water | ✗ | ✓ | ✗ |
| Nectar | ✗ | ✓ | ✓ |
| Empty | ✗ | ✗ | ✗ |

Despite extensive research on the Western honey bee, *Apis mellifera*, our understanding of how landscape composition shapes floral resources and foraging strategies in other honey bee species remains limited, even though most *Apis* diversity is concentrated in Asia [32]. Asian honey bee species like *Apis cerana* (Eastern honey bee), *Apis florea* (dwarf honey bee) and *Apis dorsata* (giant honey bee) co-occur with the introduced *A. mellifera* across overlapping habitats that range from natural forests to intensively modified agricultural and urban environments [33–36]. These species differ in body size, nesting biology, and colony organisation. Cavity-nesting *A. mellifera* and *A. cerana* inhabit tree hollows or rock crevices, while open-nesting *A. florea* and *A. dorsata* construct exposed combs on branches or rock faces [34,37,38]. Variations in such ecological, and morphological traits such as body size, likely influence how each species exploits floral resources and organises foraging labour [35,39,40]. However, comparative studies of honey bee species across landscape gradients remain scarce.

Understanding how landscape composition and species traits interact to shape foraging ecology is especially relevant in regions undergoing rapid land-use transitions, such as India, where forest conversion, agricultural intensification and urban expansion alter floral resource dynamics [41]. Comparative analyses of co-occurring *Apis* species across contrasting landscapes can therefore reveal the mechanisms by which environmental and physiological factors jointly determine foraging strategies and coexistence. Here, we investigate how landscape composition influences the foraging ecology of three co-occurring honey bee species, *A. mellifera*, *A. cerana*, and *A. florea,* across forest, agricultural, and urban landscapes in southern India. We quantified foraging preferences by characterising the nectar collected by returning foragers, compared division of foraging labour, and evaluated individual sucrose responsiveness as a behavioural correlate of the three studied honey bee species across landscapes [24,42–44].

## METHODS

### Study design and honey bee colonies

Experiments were conducted in three honey bee species: *Apis mellifera*, *Apis cerana* and *Apis florea* at a total of nine locations across three landscape types - forest, agricultural and urban in the study region, Thiruvananthapuram, Kerala, India, with two to three locations per landscape type, depending on the experiment. Each location was defined as an area of radius 1 km, chosen in such a way that it exceeded the typical foraging range of the honey bee species studied. Study locations were separated by at least 1 km distance between the circumferences to avoid overlap. Experiments were conducted in one colony per species (*A. mellifera*, *A. cerana* and *A. florea*) in each location. Colonies of *A. mellifera* and *A. cerana* were purchased from local beekeepers and established in the nine study locations. Wild *A. florea* colonies were naturally located near the study locations and were transferred to wooden boxes for experimental purposes. As the dance floor of *A. florea* is horizontal and located at the top of the hive [45–47], the boxes were left open at the top to allow unrestricted traffic of foragers. The transferred colonies were allowed to settle for two days before being used for experiments. Typically, such colonies stay in the box for several weeks. The experiments were conducted during the day between 0900 and 1500 hours local time.

Our study included two experiments (as described below).

### Experiment 1: Foraging preferences and division of foraging labour

We performed this experiment in eight locations - two forests, three agricultural and three urban locations between October 2023 and February 2024. Due to the tropical climate of the study region, temperature variations between the study periods were minimal, with day temperatures typically ranging from 26°C to 33°C throughout the year [48]. This experiment was conducted to investigate the foraging preferences and organisation of foraging labour across species and landscapes, by analysing the material brought by foragers returning to the hive (table 1).

### 1a. Analysis of crop content

Returning foragers of *A. mellifera*, *A. cerana* and *A. florea* were collected at the hive entrance in glass vials. To ensure the collection of only foragers, bees were collected when orientation flights were not observed. To prevent capturing outgoing foragers, the hive entrances were briefly closed during collection. Typically, pollen foragers are identifiable by the pollen loads on their hind legs. However, since many foragers tend to collect both pollen and nectar during their bouts [24], it was important to additionally examine their crop content to correctly determine their foraging preferences and task specialisation. Hence, the crop content of individual foragers was extracted using a protocol that allowed fast and efficient sampling of the honey bee crop through centrifugation [49]. Returning foragers were briefly immobilised on ice and individually placed upside down in a 0.5 mL microcentrifuge tube with a small hole at the tip. Each tube was then placed within a 1.5 mL microcentrifuge tube to collect the extract. The crop content of each bee was extracted immediately at the study location by centrifugation at 1300 rpm for 10 minutes using a portable minicentrifuge (Eppendorf MiniSpin^®^), which was powered by a portable power supply (UPS model: Zebronics ZEB-CU5013). The concentration of each sample was then measured using a handheld sucrose brix refractometer (Extech, Model RF10-CEI, range: 0-32 Brix %). For this refractometer model, Brix % directly represents the percentage of sucrose by weight in an aqueous solution (% w/w). The volume of each sample was also determined by pipetting. A volume of 7-10 microlitres was required for measuring concentration precisely using the refractometer. For the samples with low volumes and concentrations above the device range, a 1:10 dilution was prepared using distilled water before estimating the concentration.

#### Comparison of nectar preferences

To specifically examine how nectar preferences differ across landscapes for each species, the crop content of foragers who collected nectar (‘nectar’ and ‘pollen and nectar foragers’) was specifically analysed, excluding water foragers (crop concentration < 5 Brix %, table 1).

#### Comparison of crop volume

To examine how the volume of the crop content differs across landscapes for each species, the crop volumes of ‘nectar’, ‘pollen and nectar’, and ‘water’ (table 1) were compared. The ‘pollen and water’ foragers (table 1) were excluded from this analysis due to their small sample size.

### 1b. Foraging roles

The forager roles were determined based on the characteristics of crop content (table 1) [23–25]. The distribution of forager roles of each species was compared across landscapes to assess differences in the organisation of foraging labour.

The sample sizes for experiment 1 are summarised in table S1 and reported in the corresponding figure legends (figures 1; 2; 3).

**Figure 1:**
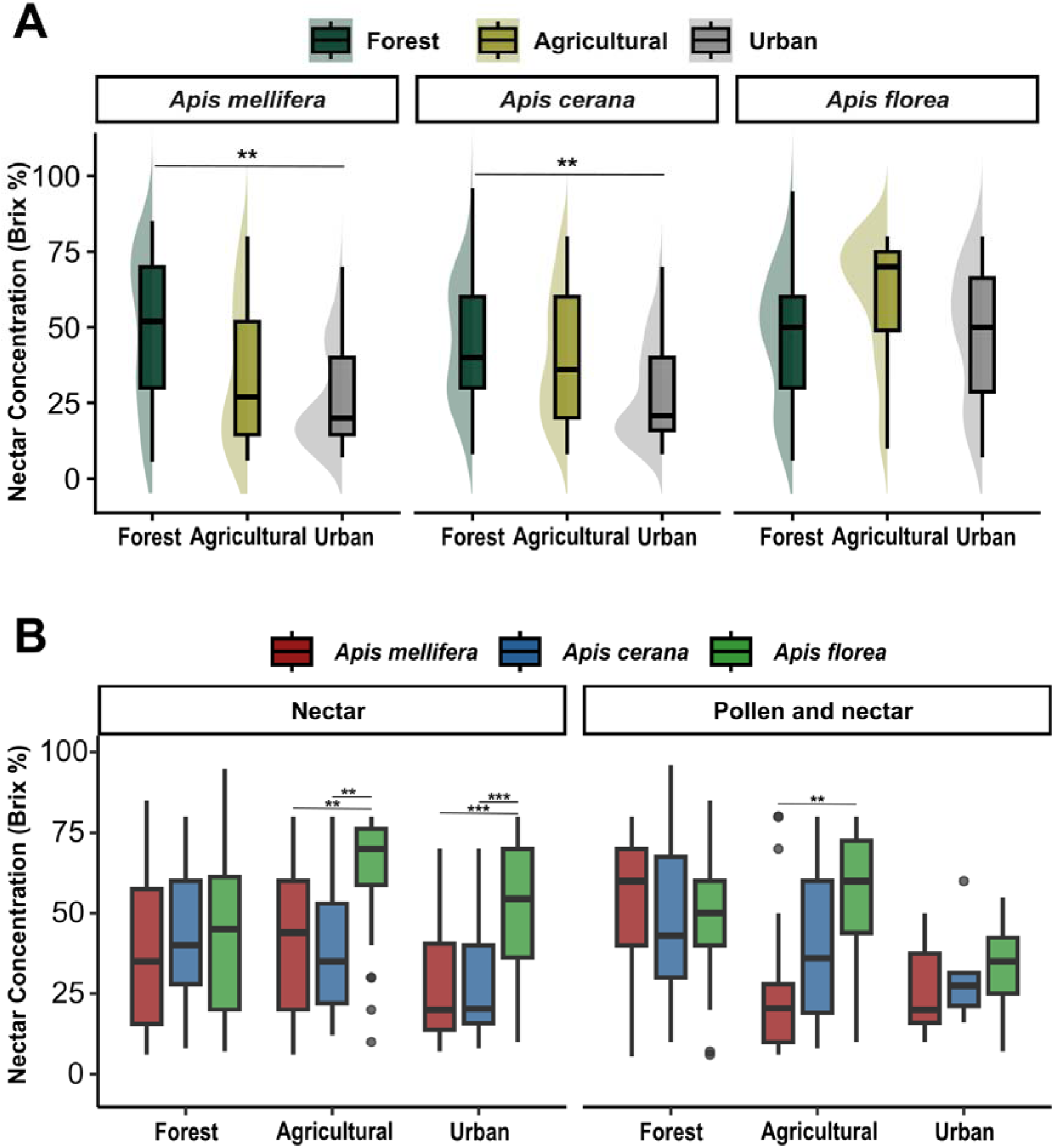
Factors influencing nectar concentration. **(A)** Concentration (Brix %) of nectar collected across forest, agricultural and urban landscapes by the three honey bee species *A. mellifera*, *A. cerana* and *A. florea*. Sample sizes for *A. mellifera*: n=47 (Forest), n=48 (Agricultural), n=55 (Urban); *A. cerana*: n=59 (Forest), n=81 (Agricultural), n=86 (Urban); *A. florea*: n=73 (Forest), n=51 (Agricultural), n=44 (Urban). Detailed sample sizes are listed in table S1. **(B)** The foraging roles, ‘Pollen’ and ‘Pollen and nectar’, are represented in the left and right columns, respectively. The asterisks (*) denote statistically significant contrasts, as determined by model-estimated marginal means from the best-fit linear mixed-effects model assessing the effects of species, landscape, and foraging role on nectar concentration (tables S4; S5). Sample sizes for Nectar (Forest): n=18 (*A. mellifera*), n=45 (*A. cerana*), n=32 (*A. florea*); Nectar (Agricultural): n=27 (*A. mellifera*), n=58 (*A. cerana*), n=32 (*A. florea*); Nectar (Urban): n=36 (*A. mellifera*), n=80 (*A. cerana*), n=36 (*A. florea*); Pollen and nectar (Forest): n=29 (*A. mellifera*), n=14 (*A. cerana*), n=41 (*A. florea*); Pollen and nectar (Agricultural): n=21 (*A. mellifera*), n=23 (*A. cerana*), n=19 (*A. florea*); Pollen and nectar (Urban): n=19 (*A. mellifera*), n=6 (*A. cerana*), n=8 (*A. florea*). Detailed sample sizes are listed in table S1.

### Experiment 2: Sucrose Responsiveness Assay

This experiment was conducted from March to June 2023 and from November 2023 to February 2024 across nine locations - three each of forest, agricultural and urban locations. We aimed to evaluate landscape- and species-level differences of honey bees in their responses to sucrose solutions. This assay was used as an indicator of foraging preferences, in addition to direct observations of crop content.

A separate set of returning foragers (different from the individuals used for Experiment 1) was collected for each species following the same procedure described for Experiment 1. Bees were classified into two roles, as either ‘pollen’ or ‘non-pollen’, based on visual examination for the presence of pollen on their legs. Subsequently, the bees were immobilised on ice and mounted in specialised tubes using tapes such that their head, antennae and mouthparts were free to move [50,51]. After this, they were placed in a humid chamber for 60 minutes for adaptation [52] before the experiment. Sucrose responsiveness of individual bees was measured by monitoring their proboscis extension responses (PER) to sucrose solutions of increasing concentrations [51]. PER is a reliable method to measure sucrose responses in hymenopterans [28,53,54]. It is the reflexive extension of the proboscis of the bee when its antennae are stimulated with a sucrose solution of sufficient concentration. The bees were stimulated successively with a series of sucrose solutions, starting with water (0%, 0.1%, 0.3%, 1%, 3%, 10% and 30% w/v sucrose). The responses were marked as a binary response of “1” or “0”, if the bee extended its proboscis or not, respectively. The gustatory response score (GRS), which is the sum of all positive responses of a bee, can range from 0 to 7 [51,55]. Between each stimulation, there was an inter-trial interval of approximately two minutes to avoid intrinsic sensitisation [56]. The sample sizes for the sucrose responsiveness assay are summarised in table S2 and reported in the corresponding figure legend (figure 4).

#### Statistical analysis

All statistical analyses and visualisations were performed using R Statistical Software version 4.3.2 [57]. The packages *AICcmodavg* [58], *car* [59], *clubSandwich* [60], *dplyr* [61], *emmeans* [62], *lme4* [63], *lmerTest* [64], *MuMIn* [65], *nnet* [66] and *ordinal* [67] were used for statistical analyses. For visualisations, the packages *gghalves* [68], *ggpattern* [69], *ggplot2* [70], *ggpubr* [71] and *tidyverse* [72] were used.We used a linear mixed effects model (*lme4* package; *lmer* function) to analyse the effects of landscape type, species and foraging role (explanatory variables) on nectar concentration and crop volume (response variables, Exp. 1a). Division of foraging labour (Exp. 1b) was analysed using a multinomial logistic regression model (*nnet* package; *multinom* function), with foraging role as the categorical response variable, and species and landscape as explanatory variables. The model estimated the probability of each foraging role across these predictors. Gustatory Response Score (GRS, Exp. 2) was analysed using a cumulative link mixed model (mixed-effects ordinal regression; *ordinal* package; *clmm* function), with landscape, species, and foraging role as explanatory variables. GRS was treated as an ordered categorical variable with values from 0 to 7. In all models, we included the relevant interactions between explanatory variables to understand the potential context-dependent effects (Refer to table S3 for an overview of the models used, including response variables, explanatory variables and random factors). All possible model combinations were compared using the Akaike Information Criterion corrected for small sample size (AICc). For models with ΔAICc < 2, the significance of predictors was checked using type II or type III Wald chi-square or Analysis of Variance tests, based on the model type. For pairwise comparisons among predictor levels in all models, we used estimated marginal means (*emmeans* package), with P-values adjusted for multiple comparisons using the Tukey method.

## RESULTS

### Sugar concentration and volume of nectar collected are shaped by landscape and species

The concentration of nectar collected varied significantly across landscapes and species (table 2; figure 1). Foragers of both *A. mellifera* and *A. cerana* collected nectar with higher concentrations in forest than in urban landscapes, whereas *A. florea* consistently foraged for high nectar concentrations across all landscape types (table S4; figure 1A). There was no significant difference between the nectar concentration collected by *A. mellifera* and *A. cerana*, in any of the landscape types (table S5). Nectar concentration collected by each species across landscapes was further influenced by the foraging role (‘nectar’ vs. ‘pollen and nectar’) (table S5; figure 1B), as indicated by the significant interactions of landscape with species and foraging role (table 2).

**Table 2:**
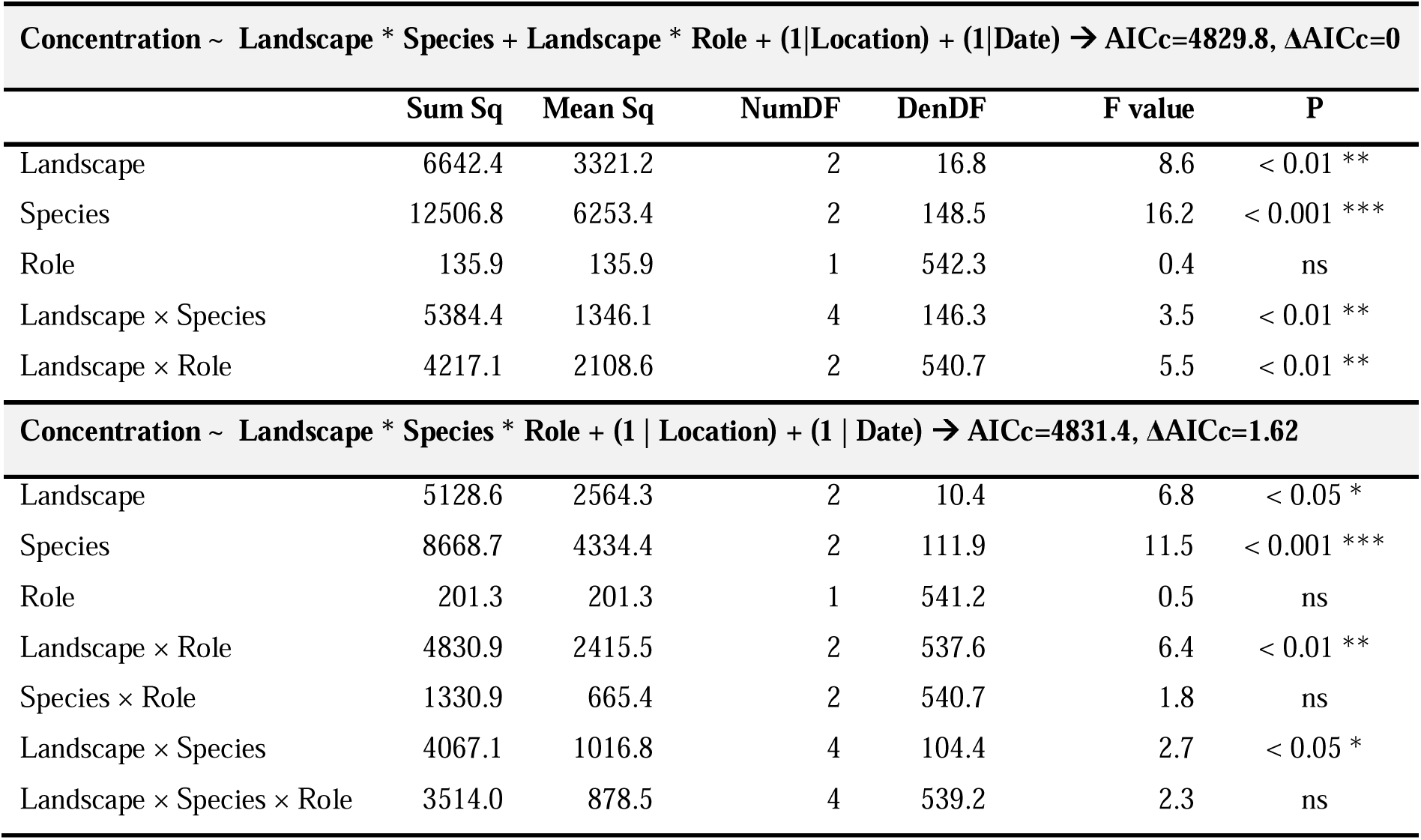
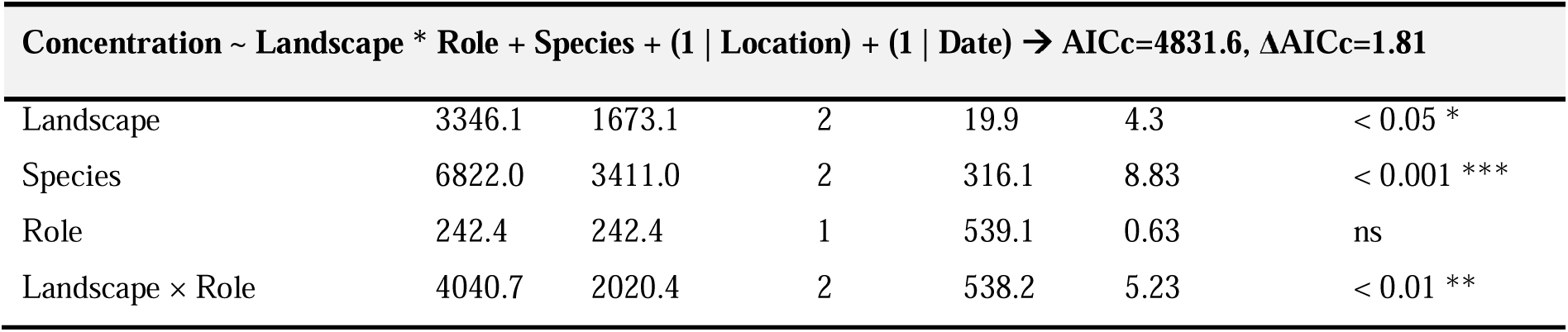
The significance of predictors (species, landscape and foraging role) on nectar concentration assessed using type III Analysis of Variance (ANOVA) with Satterthwaite’s method for the three best-fit linear mixed-effects models (ΔAICc < 2). This also includes the global model (Concentration ∼ Landscape * Species * Role + (1 | Location) + (1 | Date)). Sum Sq and Mean Sq indicate sums and means of squares. NumDF and DenDF are numerator and denominator degrees of freedom. F value is the test-statistic, and ns denotes P > 0.05.

Volume of the crop load was primarily shaped by species, likely influenced by their body size (table 3; figure 2A). This contrasts with nectar concentration, where the smallest species (*A. florea*) collected the highest concentrations. Foragers of the dwarf honey bee, *A. florea*, returned with lower crop volumes as compared to the larger *A. mellifera* and *A. cerana* foragers, irrespective of their foraging role (table S6; figure 2A). In addition to species identity, crop volume was influenced by foraging role, as indicated by the significant interactions of foraging role with species and landscape (table 3). ‘Water’ foragers of *A. mellifera* collected significantly higher volumes than ‘nectar’ foragers (table S6). Intriguingly, only in *A. florea*, nectar volume was positively correlated with nectar concentration (Spearman’s ρ = 0.33, P < 0.001), while no such correlation was detected in *A. mellifera* and *A. cerana* (figure 2B).

**Figure 2:**
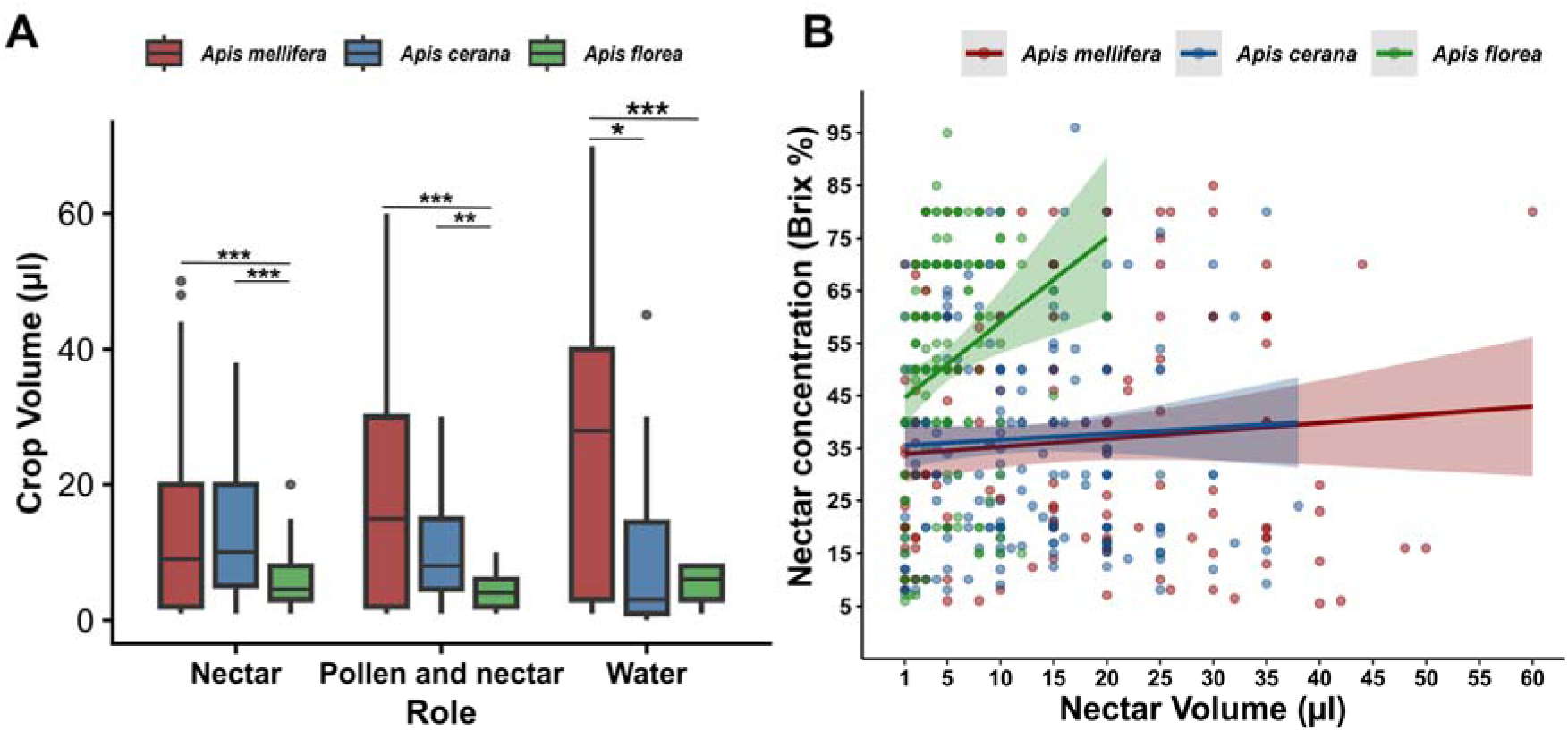
**(A)** The volume (μl) of the crop contents of different foraging roles of the honey bee species *A. mellifera*, *A. cerana* and *A. florea*. The asterisks (*) denote statistically significant contrasts, as determined by model-estimated marginal means from the best-fit linear mixed-effects model assessing the effects of species, landscape, and foraging role on crop volume (table S6). Sample sizes for Nectar: n=81 (*A. mellifera*), n=183 (*A. cerana*), n=100 (*A. florea*); Pollen and nectar: n=69 (*A. mellifera*), n=43 (*A. cerana*), n=68 (*A. florea*); Water: n=45 (*A. mellifera*), n=19 (*A. cerana*), n=9 (*A. florea*). Detailed sample sizes are listed in table S1. **(B)** shows the correlation between volume (μl) and concentration (Brix %) of nectar collected by different honey bee species. The shaded areas indicate 95% confidence intervals around the fitted regression lines.

**Table 3:** The significance of predictors (species, landscape, and foraging role) on crop volume assessed using type III Wald χ^2^ tests for the three best-fit linear mixed-effects models (ΔAICc < 2). The global model (Crop volume ∼ Landscape * Species * Role + (1 | Location) + (1 | Date)) did not emerge as a best-fit model. ns denotes P > 0.05. Df denotes degrees of freedom.

| **Crop volume ~ Role \* Landscape + Role\*Species + (1 \| Location) + (1 \| Date) → AICc=4649.3, ΔAICc=0** |  |  |  |
| --- | --- | --- | --- |
| | $\chi^2$ | Df | P |
| Role | 2.136 | 2 | ns |
| Landscape | 1.640 | 2 | ns |
| Species | 76.674 | 2 | < 0.001 *** |
| Role × Landscape | 26.164 | 4 | < 0.001 *** |
| Role × Species | 43.997 | 4 | < 0.001 *** |
| **Crop volume ~ Role \* Species + (1 \| Location) + (1 \| Date) → AICc=4650.7, ΔAICc=1.36** |  |  |  |
| Role | 1.99058 | 2 | ns |
| Species | 129.5621 | 2 | < 0.001 *** |
| Role × Species | 176.0229 | 4 | < 0.001 *** |
| **Crop volume ~ Role\*Landscape + Role\* Species + Landscape\* Species + (1 \| Location) + (1 \| Date)→**<br>**AICc=4650.9, ΔAICc=1.54** |  |  |  |
| Role | 4.163834 | 2 | ns |
| Landscape | 2.375814 | 2 | ns |
| Species | 45.4473 | 2 | < 0.001 *** |
| Role × Landscape | 27.56191 | 4 | < 0.001 *** |
| Role × Species | 69.21702 | 4 | < 0.001 *** |
| Landscape × Species | 136.8965 | 4 | < 0.001 *** |

### Landscape type influences the distribution of foraging roles for each species

Landscape type, species, and their interaction significantly contributed to the differences in the distribution of foraging roles (figure 3; table 4). Across species, the proportion of pollen foragers was generally the highest in agricultural locations, whereas forests had the most foragers collecting both pollen and nectar (table S7; figure 3). Although *A. florea* maintained similar proportions of nectar foragers across landscapes, they had an unusually high proportion of ‘empty’ (unsuccessful) foragers in urban areas, comparable to the proportion of bees collecting nectar (table S7; figure 3). On the other hand, *A. mellifera* had the fewest ‘empty’ foragers and the highest proportion of ‘pollen’ foragers in urban landscape (table S7; figure 3). Nectar foraging was notably dominant in *A. cerana*, while pollen foragers and foragers collecting both pollen and nectar were relatively less frequent (table S7; figure 3). Generally, *A. mellifera* had the highest proportion of water foragers compared to other species across landscapes (table S7; figure 3). Also, a small fraction (∼3%) of *A. mellifera* foragers returned with both pollen and water in agricultural locations (figure 3); this pattern was absent in the other species. While *A. mellifera* maintained a relatively more balanced distribution of forager types across landscapes, *A. florea* showed the most landscape-specific shifts in foraging cohort (table S7; figure 3).

**Figure 3:**
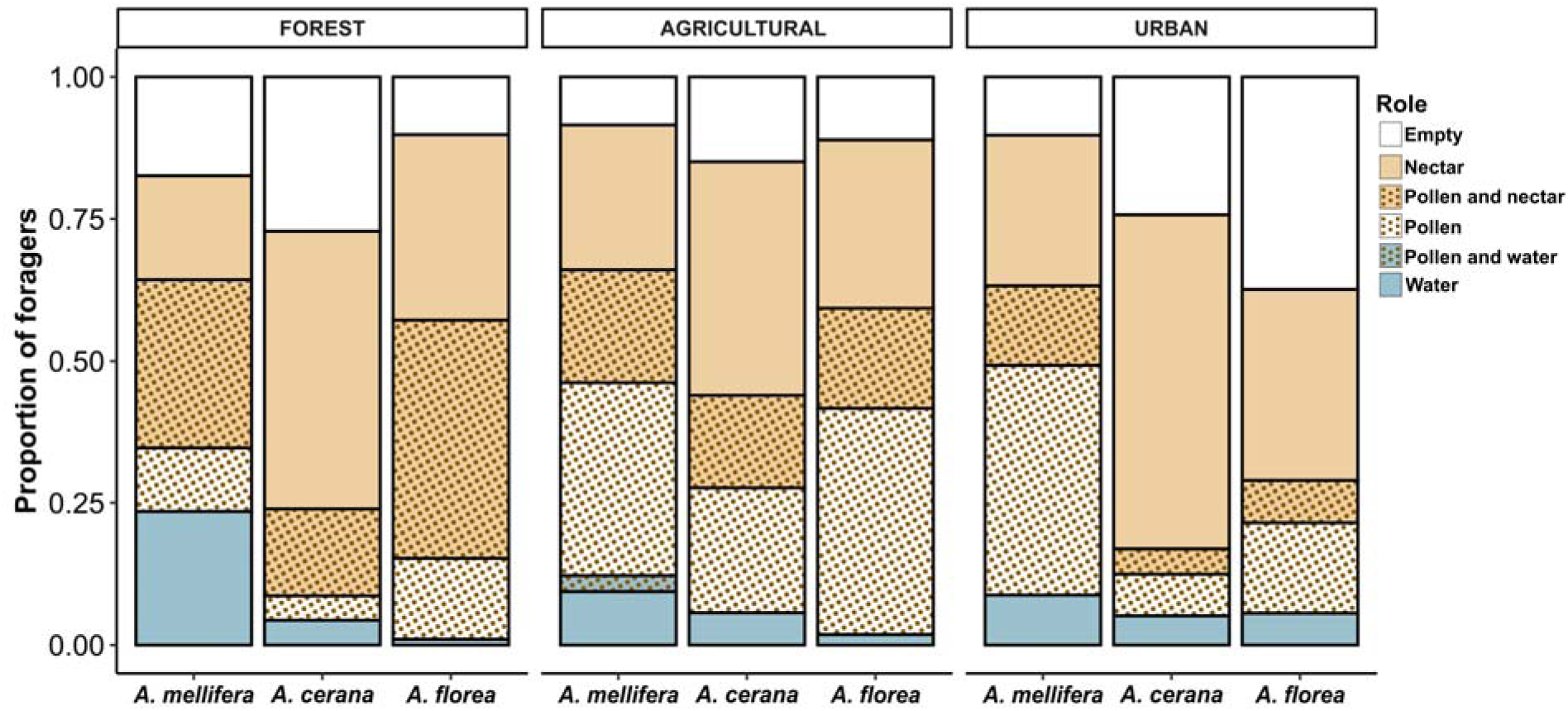
Division of foraging labour. Distributions of different foraging roles in the three honey bee species *A. mellifera*, *A. cerana* and *A. florea* across forest, agricultural and urban landscapes. The forager roles were determined based on the characteristics of the crop content and the material brought back by the bees (table 1). Significant pairwise contrasts between foraging roles are indicated in table S7. Sample sizes for Forest: n=98 (*A. mellifera*), n=92 (*A. cerana*), n=98 (*A. florea*); Agricultural: n=106 (*A. mellifera*), n=141 (*A. cerana*), n=108 (*A. florea*); Urban: n=136 (*A. mellifera*), n=136 (*A. cerana*), n=107 (*A. florea*). Detailed sample sizes, including the distribution of roles, are listed in table S1.

**Table 4:** The significance of predictors (species and landscape) on the proportion of forager roles assessed using the type II Wald χ^2^ test for the best-fit multinomial logistic regression model: Role ∼ Species * Landscape (AICc=2912.9, ΔAICc=0), which was also the global model. Df denotes degrees of freedom.

| Predictor | $\chi^2$ | Df | P |
| --- | --- | --- | --- |
| Landscape | 108.7 | 10 | < 0.001 *** |
| Species | 127.6 | 10 | < 0.001 *** |
| Landscape × Species | 53.4 | 20 | < 0.001 *** |

### Sucrose responsiveness across species and landscapes

The three honey bee species showed substantial differences in their responsiveness to sucrose (figure 4A; table 5). Across all landscapes, each species differed significantly from the others, with *A. mellifera* exhibiting the highest and *A. florea* the lowest gustatory scores (figure 4A; table S8). The effect of species on sucrose responsiveness varied between landscapes (significant interaction effect of landscape and species; table 5), while foraging role did not affect sucrose responsiveness (table 5). ‘Pollen’ foragers and ‘non-pollen’ foragers displayed similar gustatory scores.

**Figure 4:**
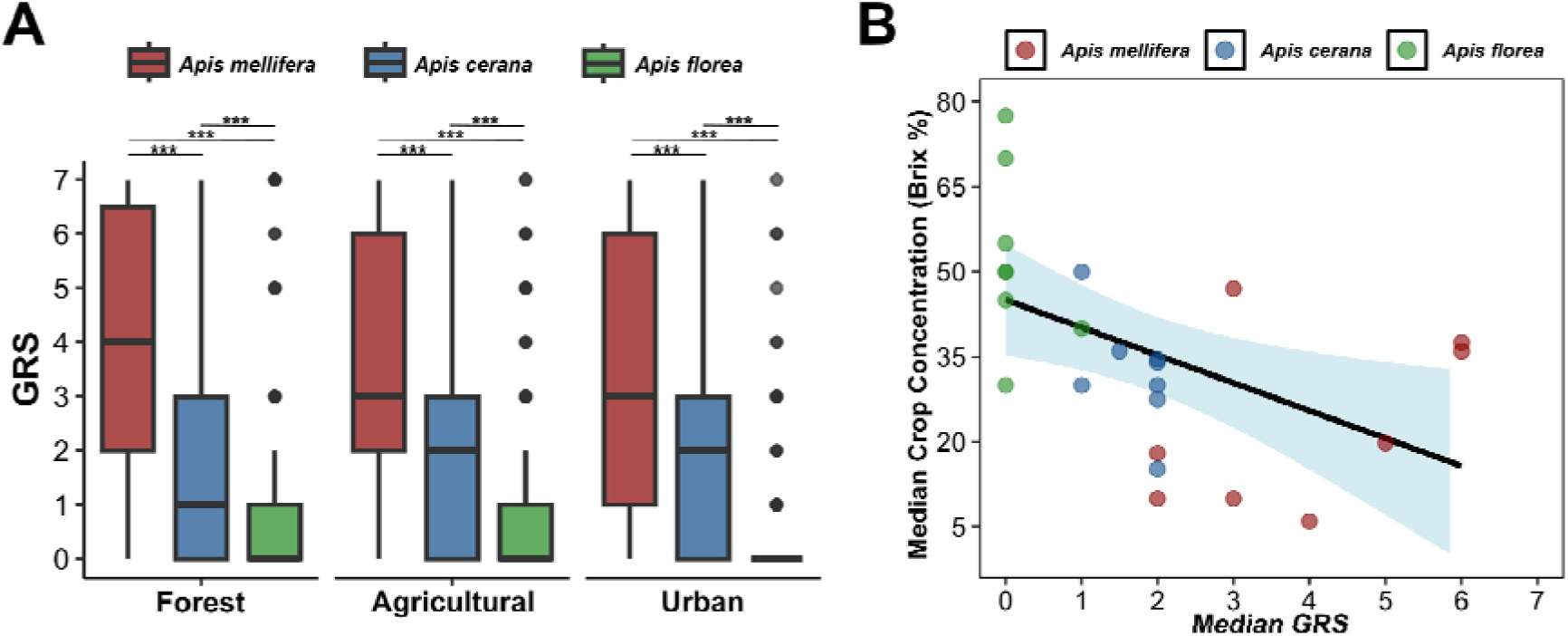
Gustatory responsiveness as an indicator of foraging preferences. **(A)** Gustatory response scores (GRS) of the three honey bee species *A. mellifera*, *A. cerana* and *A. florea* across forest, agricultural and urban landscapes. Asterisks (*) denote statistically significant differences across groups, as determined by the best-fit mixed-effects ordinal regression model (table S8). Sample sizes for Forest: n=159 (*A. mellifera*), n=178 (*A. cerana*), n=125 (*A. florea*); Agricultural: n=156 (*A. mellifera*), n=153 (*A. cerana*), n=119 (*A. florea*); Urban: n=153 (*A. mellifera*), n=149 (*A. cerana*), n=119 (*A. florea*). Detailed sample sizes are listed in table S2. **(B)** Relationship between median crop concentration and median GRS in each study location. Each point represents one species at one location (n = 24 from 8 locations). The shaded area indicates 95% confidence intervals around the fitted regression line.

**Table 5:** The significance of predictors (species and landscape) on the gustatory response scores (GRS) assessed using the type II Wald χ^2^ test for the two best-fit cumulative link mixed models (CLMM) with ΔAICc < 2. The global model (GRS ∼ Landscape * Species + Role + (1 | Date) + (1 | Location)) did not emerge as a best-fit model. Df denotes degrees of freedom and ns denotes P > 0.05.

| GRS ~ Landscape * Species + (1 \| Date) + (1 \| Location) → AICc=4463.6, ΔAICc=0 | | | |
| --- | --- | --- | --- |
| Predictor | $\chi^2$ | Df | P |
| Landscape | 0.34 | 2 | ns |
| Species | 265.86 | 2 | < 0.001 *** |
| Landscape × Species | 12.27 | 4 | < 0.05 * |
| GRS ~ Species + (1 \| Date) + (1 \| Location) → AICc=4464.2, ΔAICc=0.62 | | | |
| Species | 274.68 | 2 | < 0.001 *** |

## DISCUSSION

Our study reveals that landscape composition plays a vital role in shaping the foraging behaviour of honey bees in southern India, with distinct nectar preferences (figures 1; 2; 4A) and task allocation patterns (figure 3) across species.

### Foraging landscape shapes honey bee nectar preferences

The differences in the sugar concentration of nectar collected by species across landscapes (figure 1) suggest that landscape characteristics, such as resource availability and quality, potentially shape foraging choices. Interestingly, in forest locations, all bee species collected nectar that was similar in sugar concentration, while *A. mellifera* and *A. cerana* collected less concentrated nectar than *A. florea* in agricultural and urban areas (figure 1B). This pattern suggests differences in floral resource availability between landscapes, possibly a greater availability of flowers with higher sugar concentrations in forests, and also different preference thresholds among the three species. During the study period, we observed prominent mass-flowering tree species, such as *Terminalia paniculata*, in our forest locations, which may have served as focal foraging points for all bee species. Although several plant species were flowering, floral resource availability in the forest locations appeared to be influenced by these abundant species (*authors’ personal observation*). The forest locations seemed to have a lower overall floral diversity and understory flower cover compared to agricultural and urban locations (*authors’ personal observation*), similar to patterns reported previously [73]. This may have led to a convergence in bee foraging on a limited subset of rewarding floral resources in forest locations, resulting in the observed similarity in nectar sugar concentrations. Such limited resource diversity, with consumers focusing on available resources, may also lead to higher generalisation in pollinators [74,75]. Alternatively, urban landscapes with relatively higher floral diversity due to cultivated ornamental species and gardens [76,77] may support a greater number of coexisting pollinators, resulting in greater foraging specialisation and niche partitioning to reduce interspecific competition [74,75]. In the urban locations, we observed mass foraging of *A. florea* on a limited number of plant species, such as *Antigonon leptopus,* which provided nectar of high concentration (46.1 ± 2.8 Brix %, measured around anthesis). So, *A. florea* may have selectively targeted high-rewarding plants to maximise energetic gains, resulting in the collection of comparatively higher nectar concentrations. The other species, *A. mellifera* and *A. cerana,* may have exploited a wider range of nectar sources with lower sugar concentration but higher abundance or accessibility, which may still be energetically profitable given their ability to collect larger volumes due to their larger body size (figure 2A) [78,79].

The significant interaction between landscape and foraging role emphasises context-dependent foraging preferences (table 2). In agricultural locations, *A. mellifera* ‘nectar’ foragers collected nectar with higher concentrations than ‘pollen and nectar’ foragers, whereas in forests, the opposite pattern was observed with ‘nectar’ foragers collecting lower concentrations than ‘pollen and nectar’ foragers (table S5; figure 1B). In urban landscapes, *A. florea* ‘nectar’ foragers collected nectar with higher concentrations than those of other species, whereas ‘pollen and nectar’ foragers across species collected similar concentrations (table S5; figure 1B). Together, these patterns suggest flexibility in honey bee foraging behaviour across landscapes. Differences in resource profitability among landscapes may underlie the observed variation in the nectar concentrations collected by specialised and mixed nectar foragers, and may reflect adjustments in foraging strategies to maximise colony efficiency [16].

### Species-specific foraging strategies

Intriguingly, each species responded differently to similar landscape conditions, indicating distinct foraging strategies and habitat suitability. While *A. florea* consistently preferred high-concentration nectar, *A. mellifera* and *A. cerana* seemed to be more flexible in terms of foraging choices.

Species-level differences in nectar preferences may potentially be shaped by body size, colony size, physiology and nesting biology [35,78,79]. The medium-sized cavity-nesting bees (*A. mellifera* and *A. cerana*) have greater flight speeds, thoracic temperatures, and metabolic rates, and make more foraging trips per day than the large (*A. dorsata*) and small-sized (*A. florea*) open-nesters [35,80]. However, the cavity-nesting species also have shorter lifespans than open-nesting species [35]. The open-nesting species have longer lifespans and allocate a substantial proportion of their adult workforce for thermoregulation and colony defence [47,81,82], limiting the number of brood cells, foragers and foraging trips [35]. Maximising returns from each trip becomes critical for *A. florea*, given its smaller colony size, along with a limited number of adults available for foraging compared to the other two species [35,83].

The distribution of nectar concentrations across landscapes (figure 1A) indicates that the nectar collected by *A. florea* was typically more concentrated than that collected by *A. mellifera* and *A. cerana* in agricultural and urban landscapes (figure 1; table S5). This consistent preference of *A. florea* for higher nectar concentrations likely reflects a strategy to maximise energy efficiency, shaped by their distinct physiological traits. The small size and low thoracic temperature gradient of *A. florea* [37] may favour foraging at warmer times of the day [33], when many flowers have high-concentration nectar [84,85], increasing the energy carried per unit volume [86].

Although *A. florea* generally collects smaller nectar volumes than the cavity-nesters due to small body size, we found a positive correlation between nectar volume and concentration within this species (figure 2B). This suggests that when *A. florea* encounters very high-concentration nectar, it tends to collect larger crop loads, thereby maximising the foraging efficiency by carrying maximum provisions with its limited crop volume [86]. The possibility of an increase in nectar concentration via evaporative cooling is unlikely for *A. florea* due to its fast cooling rate due to small size [37]. In contrast, the absence of correlation in *A. mellifera* and *A. cerana* (figure 2B) indicates a different strategy, potentially shaped by their greater colony demands [35,40] or different nectar preferences [87]. Previous field observation studies on flower visitation patterns of *A. mellifera* and *A. cerana* show that both species often bias their foraging towards flowers offering ∼30% w/w nectar concentration, even when richer (>50% w/w) nectar sources are available [87]. This aligns with our observation that their collected nectar was generally less concentrated than that of *A. florea*. Moreover, stronger preference in *A. mellifera* for flowers with larger nectar volume compared to *A. cerana* [87] resonates with our interpretation that its foraging strategy may be more volume-oriented, maximising total intake per trip rather than absolute nectar concentration. In addition, low-concentration and less viscous nectar enables faster fluid uptake, which is advantageous when collecting larger volumes [79]. Alternatively, this might also be linked to the increased demands of water collection in *A. mellifera,* since low-concentration nectar may provide more water that can be used for hive cooling and thermoregulation.

The differences in nectar preferences across species are further emphasised by their sucrose responsiveness (figure 4A), suggesting that species-specific traits and foraging preferences may be closely linked to their responsiveness to sugar rewards. The high sucrose responsiveness in *A. mellifera* and *A. cerana* (figure 4A) aligns with their collection of comparatively lower-concentration nectar and is also consistent with previous comparative studies [88]. Conversely, the low sucrose responsiveness of *A. florea* across landscapes (figure 4A) reflects its consistent preference for high-concentration nectar (figures 1; 2B). Notably, the median concentration of the nectar brought back by *A. florea* was always higher (∼50 Brix %) than the maximum sucrose concentration used in the sucrose responsiveness assay (30% w/v), which could also contribute to the low responsiveness of *A. florea* in the assay. We further observed a strong negative correlation between median gustatory response score (GRS) and crop concentration (Spearman’s ρ = −0.61, P < 0.01; figure 4B), suggesting that bees with higher gustatory responsiveness tend to collect resources with lower sugar concentrations. The complementary results from both direct measurements of nectar concentrations brought by returning foragers and the indirect sucrose responsiveness assay further highlight that the assay provides a reliable proxy for estimating nectar preferences in bees. Similarly, the significant species × landscape interaction in sucrose responsiveness (table 5) hints that the resource environment may modulate species-level differences in GRS, even though landscape was not a significant main effect influencing sucrose responsiveness. Our results indicate that foraging strategies of honey bees are likely influenced by interspecific differences in physiology, body size, and nesting behaviour, and reveal possible drivers of resource partitioning, minimising competition and fostering coexistence.

### Division of foraging labour across landscapes and species

Our study is the first to reveal clear landscape- and species-dependent patterns in the task specialisation in honey bee foraging. The prominent landscape influence suggests that resource availability and foraging environment can substantially modify colony-level foraging strategies and thereby affect the composition of the foraging cohort. The increased pollen foraging observed across all species in agricultural locations likely reflects the greater availability of preferred pollen resources in these landscapes. Notably, coconut (*Cocos nucifera*) and *Mimosa pudica*, which are excellent pollen sources for bees [89], were abundant in our agricultural locations (*authors’ personal observation*). We observed substantial bee visitation on many nectarless flowers (flowers producing only pollen), such as *Mimosa pudica*, in agricultural locations during the study period (*authors’ personal observation*). On the contrary, the large proportion of bees collecting both pollen and nectar observed in forests (figure 3) may reflect a foraging strategy to maximise overall nectar intake in forests, aligning with the observation that all species collected high-sugar nectar in forest locations (figure 1). Previous work has shown that ‘pollen and nectar’ and ‘nectar’ foragers have similar sucrose response thresholds [25], which may facilitate the collection of both resources. Alternatively, the increased generalist foragers (‘pollen and nectar’) in forests could also be an artefact of the generally low understory flower cover and floral diversity in this landscape (*authors’ personal observation*). This aligns with earlier studies showing that foragers collecting both resources (‘pollen and nectar’) were found to be higher in number when pollen stores were low [90]. We observe this pattern consistently across species in our study, with pure ‘pollen’ foragers being low in forests across species (figure 3).

Species-level differences in forager composition further highlight contrasting foraging strategies. The greatest landscape-influenced changes in forager composition were exhibited by *A. florea*, highlighting a flexible foraging cohort based on local resource availability to optimise its foraging efficiency (figure 3). However, in urban areas, *A. florea* had similar proportions of ‘nectar’ and ‘empty’ foragers (figure 3), indicating a higher rate of unsuccessful foraging in this landscape. Crop sugar concentration patterns (figure 1) revealed that in urban locations, *A. florea* selectively returned with high-concentration nectar, even when other species returned with comparatively lower-concentration nectar in this landscape (figure 1). This behaviour suggests a limited ability of this species in exploiting lower-quality nectar sources, even during unsuccessful foraging bouts, but rather selectively foraging on the few high-rewarding plants. This could be a constraint of their sucrose thresholds as indicated by their lower sucrose responsiveness (figure 4A). Most *A. florea* individuals remained unresponsive to sucrose concentrations up to 30% w/v (figure 4A). In contrast, the fewer empty foragers and the large proportion of pollen foragers of *A. mellifera* in urban locations suggest greater adaptability in utilising available resources, despite low sugar concentrations (figure 1), further supported by their large variation in gustatory response scores (figure 4A). The predominance of nectar foragers in *A. cerana* across landscapes is consistent with the year-round dominance of this forager group over other roles in this species [33,91]. This may further reflect a colony-level focus on meeting carbohydrate demands, particularly if pollen stores are sufficient and brood rearing was limited during the experimental months. It might also display a genetically encoded preference for nectar collection similar to the strain-level differences in pollen-hoarding behaviour reported in *A. mellifera* [92].

Further, *A. mellifera* showed the highest proportion of water foragers among all species, particularly in forest locations. This likely reflects its greater colony-level demands for thermoregulation [93–95], especially crucial in warmer tropical environments, for a species which has been adapted to temperate climates [96]. The volume of water collected by *A. mellifera* was also significantly higher than that of nectar (table S6; figure 2A), emphasising the importance of water foraging for their physiological needs [97]. The preference for lower-concentration nectar by *A. mellifera* in agricultural and urban landscapes could also be related to its water demands, since low-concentration nectar provides more water that can be used for thermoregulation.

The relatively balanced distribution of foraging roles in *A. mellifera* across landscapes (figure 3) may point to a generalist foraging strategy, which, together with its large honey production, may have facilitated its worldwide distribution through beekeeping [98]. This foraging flexibility contributes to its resilience and ecological success worldwide [99]. In contrast, *A. florea* exhibits a more specialised foraging strategy, consistently targeting high-concentration nectar and often returning empty rather than collecting lower-quality resources (figure 1; 3). While such selectivity may maximise its foraging efficiency in resource-rich environments, it could reduce resilience in resource-poor landscapes. Based on our observations that the *A. florea* colonies were found more frequently in the study region during the dry months, from December to March (*supported by year-round observations of authors*), we hypothesise that the colonies move into the landscape during periods of preferred nectar availability, and possibly leave when these resources decline. Nevertheless, *A. florea* has demonstrated ecological adaptability and invasive potential and has been expanding westwards, through both natural and accidental introduction [100,101], indicating its ability to successfully establish in new habitats.

## Conclusions

This study advances our understanding of how both landscape context and species-level traits shape honey bee foraging behaviour, particularly among the understudied Asian species. We present comprehensive evidence of diverse foraging preferences linked to foraging landscape, physiology and nesting behaviour, and report for the first time a comparison of the organisation of foraging labour in honey bee species beyond *A. mellifera*. We also show that the sucrose responsiveness assay reliably reflects foraging preferences, complementing direct measurements of nectar concentrations collected by foragers. Together, our findings support the broader view that foraging preferences of bees are flexible, allowing them to adjust their foraging labour under fluctuating ecological conditions [27,55,102]. Species-specific foraging strategies of co-occurring *Apis* species likely promote niche differentiation mechanisms and minimise competition under changing environmental and land-use conditions and also offer insights into the adaptation of the Western honey bee (*A. mellifera*) to tropical environments. With a global decline in bee populations, often driven by habitat loss and fragmentation [103,104], it is critical to address the effects of land-use changes on honey bee foraging behaviour to understand how they respond to and cope with anthropogenic changes in the foraging landscape. Comprehending these relationships is critical not only to predict the resilience of bee populations in changing landscapes and habitat suitability but also to extend this knowledge to design bee-friendly habitats. With the insights from honey bees, we can derive broader strategies for wild bee conservation and landscape management to sustain pollinator diversity and ecosystem services.

## Data accessibility

The data related to this study will be provided upon request from the corresponding authors.

## Authors’ contributions

**G.A.J.:** Methodology, Validation, Formal analysis, Investigation, Data curation, Writing – original draft, Writing – review & editing, Visualisation; **M.R.:** Methodology, Writing – review & editing; **S.B.:** Methodology, Writing – review & editing; **A.A.:** Methodology, Writing – review & editing; **H.S.:** Project administration, Resources, Supervision, Writing – review & editing; **I.S.-D.:** Funding acquisition, Project administration, Resources, Supervision, Writing – review & editing; **R.S.:** Conceptualisation, Methodology, Funding acquisition, Project administration, Resources, Supervision, Writing – review & editing.

## Acknowledgements

We thank IISER Thiruvananthapuram and the Behavioural and Evolutionary Ecology Laboratory (BEE Lab) for providing the infrastructure and logistic support necessary to conduct the experiments. We thank the Kerala Forest Department for granting permission to conduct fieldwork at our forest locations in the Peppara Wildlife Sanctuary (permit number: KFDHQ-2542/2022-CWW/WL-10). We also thank the landowners of our agricultural and urban locations for allowing the fieldwork. We acknowledge our technical assistants, Azmi Sheriff and Sivachand M. P., for support during the experiments. We thank Dr. Benjamin Rutschmann for his guidance in standardising the establishment of open-nesting honey bee colonies used in this study.

