## Supplementary Material for "Human landscape modification shapes foraging preferences and sucrose responsiveness of honey bees in Asia"

**Affiliations:**

### SUPPLEMENTARY MATERIAL

**Table S1:** Sample size and foraging role distribution for the dataset from crop content analysis (experiment 1).

| Landscape | Species | Sample size | Role | Distribution |
|---|---|---|---|---|
| Forest | *Apis mellifera* | 98 | Empty | 17 |
| | | | Nectar | 18 |
| | | | Pollen | 11 |
| | | | Pollen and nectar | 29 |
| | | | Water | 23 |
| | *Apis cerana* | 92 | Empty | 25 |
| | | | Nectar | 45 |
| | | | Pollen | 4 |
| | | | Pollen and nectar | 14 |
| | | | Water | 4 |
| | *Apis florea* | 98 | Empty | 10 |
| | | | Nectar | 32 |
| | | | Pollen | 14 |
| | | | Pollen and nectar | 41 |
| | | | Water | 1 |
| Agricultural | *Apis mellifera* | 106 | Empty | 9 |
| | | | Nectar | 27 |
| | | | Pollen | 36 |
| | | | Pollen and nectar | 21 |
| | | | Pollen and water | 3 |
| | | | Water | 10 |
| | *Apis cerana* | 141 | Empty | 21 |
| | | | Nectar | 58 |
| | | | Pollen | 31 |
| | | | Pollen and nectar | 23 |
| | | | Water | 8 |
| | *Apis florea* | 108 | Empty | 12 |
| | | | Nectar | 32 |
| | | | Pollen | 43 |
| | | | Pollen and nectar | 19 |
| | | | Water | 2 |
| Urban | *Apis mellifera* | 136 | Empty | 14 |
| | | | Nectar | 36 |
| | | | Pollen | 55 |
| | | | Pollen and nectar | 19 |
| | | | Water | 12 |
| | *Apis cerana* | 136 | Empty | 33 |
| | | | Nectar | 80 |
| | | | Pollen | 10 |
| | | | Pollen and nectar | 6 |
| | | | Water | 7 |
| | *Apis florea* | 107 | Empty | 40 |

| | | |
|---|---|---|
| | Nectar | 36 |
| | Pollen | 17 |
| | Pollen and nectar | 8 |
| | Water | 6 |

**Table S2:** Sample sizes for sucrose responsiveness assay (experiment 2).

| Landscape | Species | Role | Sample size |
|---|---|---|---|
| Forest | *Apis mellifera* | Non-pollen | 73 |
| | | Pollen | 86 |
| | *Apis cerana* | Non-pollen | 88 |
| | | Pollen | 90 |
| | *Apis florea* | Non-pollen | 86 |
| | | Pollen | 39 |
| Agricultural | *Apis mellifera* | Non-pollen | 42 |
| | | Pollen | 114 |
| | *Apis cerana* | Non-pollen | 83 |
| | | Pollen | 70 |
| | *Apis florea* | Non-pollen | 49 |
| | | Pollen | 70 |
| Urban | *Apis mellifera* | Non-pollen | 67 |
| | | Pollen | 86 |
| | *Apis cerana* | Non-pollen | 83 |
| | | Pollen | 66 |
| | *Apis florea* | Non-pollen | 77 |
| | | Pollen | 42 |

**Table S3**: Overview of the statistical models used in the study, showing response variables, explanatory variables, and random factors.

| Exp. | Model | Response variable | Explanatory variables and interactions (with levels) | Random factors |
|---|---|---|---|---|
| 1a | Linear mixed-effects model (*lme4* package) | Nectar concentration,<br>Crop volume | **Species** (*A. mellifera, A. cerana, A. florea*)<br>**Landscape** (Forest, Agricultural, Urban)<br>**Role** (Nectar, Pollen & nectar)<br>**Species × Landscape**<br>**Role × Landscape**<br>**Role × Landscape × Species** | Location, Date |
| 1b | Multinomial logistic regression (*nnet* package) | Foraging role | **Species** (*A. mellifera, A. cerana, A. florea*)<br>**Landscape** (Forest, Agricultural, Urban)<br>**Species × Landscape** | Not included |
| 2 | Cumulative link mixed model (*ordinal* package) | Gustatory Response Score | **Species** (*A. mellifera, A. cerana, A. florea*)<br>**Landscape** (Forest, Agricultural, Urban)<br>**Role** (Pollen, Non-pollen)<br>**Species × Landscape** | Location, Date |

**Table S4**: Pairwise comparisons of nectar concentrations collected from each landscape by each species (landscape contrasts), based on estimated marginal means from the best-fit linear mixed-effects model: *Concentration ~ Role × Landscape + Species × Landscape + (1 | Location) + (1 | Date).* P-values are adjusted for multiple comparisons using Tukey's method. 'ns' denotes P > 0.05. SE denotes standard error, df denotes degrees of freedom, and t-ratio is the test statistic (ratio between estimate and standard error).

| Contrast | Estimate | SE | df | t-ratio | P-value |
|---|---|---|---|---|---|
| **Species = *Apis mellifera*** | | | | | |
| Forest - Agricultural | 12.19194 | 5.995891 | 51.35 | 2.033 | ns |
| Forest - Urban | 20.43065 | 6.033498 | 48.85 | 3.386 | < 0.01 ** |
| Agricultural - Urban | 8.23871 | 6.033028 | 58.29 | 1.366 | ns |
| **Species = *Apis cerana*** | | | | | |
| Forest - Agricultural | 7.47211 | 5.703992 | 34.90 | 1.310 | ns |
| Forest - Urban | 19.243450 | 5.868400 | 40.89 | 3.279 | < 0.01 ** |
| Agricultural - Urban | 11.771341 | 5.765673 | 45.68 | 2.042 | ns |
| **Species = *Apis florea*** | | | | | |
| Forest - Agricultural | -9.446833 | 5.895358 | 31.97 | -1.602 | ns |
| Forest - Urban | 3.694315 | 5.952045 | 38.76 | 0.621 | ns |
| Agricultural - Urban | 13.141149 | 6.350660 | 47.27 | 2.069 | ns |

**Table S5:** Pairwise comparisons of nectar concentrations collected by different species within each landscape and foraging role (species contrasts), and among foraging roles (role contrasts), based on estimated marginal means from the best-fit linear mixed-effects model including the Role × Species interaction (ΔAICc = 1.62): *Concentration ~ Role × Species × Landscape + (1 | Location) + (1 | Date).* P-values are adjusted for multiple comparisons using Tukey's method. 'ns' denotes P > 0.05. SE denotes standard error, df denotes degrees of freedom, and t-ratio is the test statistic (ratio between estimate and standard error).

| Contrast | Estimate | SE | df | t-ratio | P-value |
|---|---|---|---|---|---|
| **SPECIES CONTRAST** | | | | | |
| **ROLE = Nectar, LANDSCAPE = Forest** | | | | | |
| *A. mellifera - A. cerana* | -6.14266 | 5.558703 | 552.3681 | -1.10505 | ns |
| *A. mellifera - A. florea* | -8.48368 | 6.393834 | 439.0018 | -1.32685 | ns |
| *A. cerana - A. florea* | -2.34102 | 5.377145 | 277.9243 | -0.43537 | ns |
| **ROLE = Nectar, LANDSCAPE = Agricultural** | | | | | |
| *A. mellifera - A. cerana* | 0.940305 | 4.612882 | 543.3423 | 0.203843 | ns |
| *A. mellifera - A. florea* | -20.3431 | 6.585729 | 52.39737 | -3.08897 | < 0.01 ** |
| *A. cerana - A. florea* | -21.2834 | 6.084101 | 30.16541 | -3.4982 | < 0.01 ** |
| **ROLE = Nectar, LANDSCAPE = Urban** | | | | | |
| *A. mellifera - A. cerana* | -0.69712 | 4.013168 | 549.4798 | -0.17371 | ns |
| *A. mellifera - A. florea* | -22.7286 | 5.91877 | 95.05551 | -3.84009 | < 0.001 ** |
| *A. cerana - A. florea* | -22.0315 | 5.263475 | 68.94331 | -4.18573 | < 0.001 ** |

| | | | | | |
|---|---|---|---|---|---|
| **ROLE = Pollen & Nectar, LANDSCAPE = Forest** | | | | | |
| *A. mellifera - A. cerana* | 9.648206 | 6.55091 | 555.5985 | 1.472804 | ns |
| *A. mellifera - A. florea* | 4.016646 | 5.65583 | 313.2795 | 0.710178 | ns |
| *A. cerana - A. florea* | -5.63156 | 6.545528 | 467.1444 | -0.86037 | ns |
| **ROLE = Pollen & Nectar, LANDSCAPE = Agricultural** | | | | | |
| *A. mellifera - A. cerana* | -11.2405 | 6.026885 | 550.0091 | -1.86506 | ns |
| *A. mellifera - A. florea* | -26.9053 | 7.517549 | 100.2632 | -3.579 | < 0.01 ** |
| *A. cerana - A. florea* | -15.6648 | 7.524325 | 79.9024 | -2.08189 | ns |
| **ROLE = Pollen & Nectar, LANDSCAPE = Urban** | | | | | |
| *A. mellifera - A. cerana* | -5.58797 | 9.504003 | 559.5579 | -0.58796 | ns |
| *A. mellifera - A. florea* | -4.87482 | 9.090239 | 364.4466 | -0.53627 | ns |
| *A. cerana - A. florea* | 0.713143 | 11.0469 | 551.1936 | 0.064556 | ns |
| **ROLE CONTRAST** (only significant contrasts with P-value < 0.05 are shown) | | | | | |
| **Landscape = Forest, Species = *A. mellifera*** | | | | | |
| Nectar - Pollen & nectar | -17.581626 | 6.181477 | 557.42 | -2.844 | < 0.01 ** |
| **Landscape = Agricultural, Species = *A. mellifera*** | | | | | |
| Nectar - Pollen & nectar | 13.097582 | 5.809197 | 548.82 | 2.255 | < 0.05 * |
| **Landscape = Urban, Species = *A. florea*** | | | | | |
| Nectar - Pollen & nectar | 19.335933 | 7.935403 | 563.10 | 2.437 | < 0.05 * |

**Table S6:** Pairwise comparisons of crop volumes collected by different species and foraging roles (species and role contrasts), based on estimated marginal means from the best-fit linear mixed-effects model: *Crop volume ~ Role × Landscape + Role × Species + (1 | Location) + (1 | Date).* As residual diagnostics indicated heteroscedasticity (funnel-shaped pattern), heteroscedasticity-robust standard errors were computed using the CR2 cluster-robust estimator clustered by Location (*clubSandwich* package), ensuring that significance tests are robust to unequal residual variance across groups. P-values are adjusted for multiple comparisons using Tukey's method. Only significant pairwise contrasts (P < 0.05) are shown. SE denotes standard error, df denotes degrees of freedom, and t-ratio is the test statistic (ratio between estimate and standard error).

| Contrast | Estimate | SE | df | t-ratio | P-value |
|---|---|---|---|---|---|
| **SPECIES CONTRAST** | | | | | |
| **ROLE = Nectar** | | | | | |
| *A. mellifera - A. florea* | 7.215943 | 1.808841 | 615.5626 | 3.989263 | < 0.001 *** |
| *A. cerana - A. florea* | 6.527757 | 0.789689 | 615.1555 | 8.266242 | < 0.001 *** |
| **ROLE = Pollen & nectar** | | | | | |
| *A. mellifera - A. florea* | 12.91299 | 2.465413 | 609.0409 | 5.237657 | < 0.001 *** |
| *A. cerana - A. florea* | 5.582612 | 1.76028 | 616.177 | 3.171433 | < 0.01 ** |
| **ROLE = Water** | | | | | |
| *A. mellifera - A. cerana* | 13.49373 | 4.72885 | 615.6671 | 2.85349 | < 0.05 * |
| *A. mellifera - A. florea* | 18.3176 | 3.68067 | 610.9436 | 4.976703 | < 0.001 *** |
| **ROLE CONTRAST** | | | | | |
| **SPECIES = *A. mellifera*** | | | | | |
| Nectar - Water | -10.8473 | 3.907648 | 616.9712 | -2.77591 | < 0.05 * |

**Table S7:** Significant pairwise contrasts based on differences in predicted probabilities of forager roles estimated from the best-fit multinomial logistic regression model (*Role ~ Species × Landscape*). Contrasts include comparisons among landscapes (overall and within each species), among roles, and among species. Only significant pairwise comparisons are shown. P-values are adjusted using Tukey's method. SE denotes standard error, df denotes degrees of freedom, and t-ratio is the test statistic (ratio between estimate and standard error). Pairwise contrasts involving the 'pollen and water' role were excluded from interpretation (also not shown in the table) as this role was not present for all species across landscapes.

| Contrast | Estimate | SE | df | t-ratio | P-value |
|---|---|---|---|---|---|
| **CONTRAST: OVERALL LANDSCAPE** | | | | | |
| Agricultural – Urban (**Empty**) | -0.12482 | 0.027415 | 45 | -4.55301 | < 0.001 *** |
| Forest – Agricultural (**Pollen & nectar**) | 0.109768 | 0.033033 | 45 | 3.322979 | < 0.01 ** |
| Forest – Urban (**Pollen & nectar**) | 0.202625 | 0.029537 | 45 | 6.860112 | < 0.001 *** |
| Agricultural – Urban (**Pollen & nectar**) | 0.092857 | 0.025055 | 45 | 3.706179 | < 0.01 ** |
| Forest – Agricultural (**Pollen**) | -0.21968 | 0.030312 | 45 | -7.24748 | < 0.001 *** |
| Forest - Urban (**Pollen**) | -0.11275 | 0.026329 | 45 | -4.2823 | < 0.001 *** |
| Agricultural – Urban (**Pollen**) | 0.106935 | 0.031749 | 45 | 3.368136 | < 0.01 ** |
| **CONTRAST: LANDSCAPE (within each species)** | | | | | |
| **Species =** ***A. mellifera*** | | | | | |
| Forest – Urban (**Pollen & nectar**) | 0.156216 | 0.054861 | 45 | 2.847479 | < 0.05 * |
| Forest – Agricultural (**Pollen**) | -0.22739 | 0.05597 | 45 | -4.0627 | < 0.001 *** |
| Forest - Urban (**Pollen**) | -0.29218 | 0.0528 | 45 | -5.53368 | < 0.001 *** |
| Forest – Agricultural (**Water**) | 0.140367 | 0.05137 | 45 | 2.732472 | < 0.05 * |
| Forest – Urban (**Water**) | 0.146469 | 0.049238 | 45 | 2.974725 | < 0.05 * |
| **Species =** ***A. cerana*** | | | | | |
| Agricultural – Urban (**Nectar)** | -0.17689 | 0.059146 | 45 | -2.99072 | < 0.05 * |
| Forest – Urban (**Pollen & nectar**) | 0.108056 | 0.041382 | 45 | 2.611199 | < 0.05 * |
| Agricultural – Urban (**Pollen & nectar**) | 0.119003 | 0.035753 | 45 | 3.328499 | < 0.01 ** |
| Forest - Agricultural (**Pollen**) | -0.17638 | 0.040847 | 45 | -4.31799 | < 0.001 *** |
| Agricultural - Urban (**Pollen**) | 0.146323 | 0.041441 | 45 | 3.530914 | < 0.01 ** |
| **Species =** ***A. florea*** | | | | | |
| Forest – Urban (**Empty**) | -0.27178 | 0.055882 | 45 | -4.86343 | < 0.001 *** |
| Agricultural – Urban (**Empty**) | -0.26272 | 0.055697 | 45 | -4.71686 | < 0.001 *** |
| Forest – Agricultural (**Pollen & nectar**) | 0.242446 | 0.06185 | 45 | 3.919933 | < 0.001 *** |
| Forest – Urban (**Pollen & nectar**) | 0.343602 | 0.055942 | 45 | 6.142095 | < 0.001 *** |
| Forest - Agricultural (**Pollen**) | -0.25529 | 0.058892 | 45 | -4.33492 | < 0.001 *** |
| Agricultural - Urban (**Pollen**) | 0.239269 | 0.058887 | 45 | 4.063184 | < 0.001 *** |
| **CONTRAST: ROLE (within each species)** | | | | | |
| **Species =** ***A. mellifera,*** **Landscape = Agricultural** | | | | | |
| Empty - Nectar | -0.16981 | 0.054147 | 45 | -3.13617 | < 0.05 * |
| Empty - Pollen | -0.25473 | 0.058248 | 45 | -4.37313 | < 0.001 *** |
| Pollen - Water | 0.245291 | 0.059384 | 45 | 4.130625 | < 0.01 ** |
| **Species =** ***A. mellifera,*** **Landscape = Urban** | | | | | |
| Empty - Nectar | -0.16175 | 0.050109 | 45 | -3.22805 | < 0.05 * |
| Empty - Pollen | -0.30147 | 0.055338 | 45 | -5.44778 | < 0.001 *** |

| | | | | | |
|---|---|---|---|---|---|
| Nectar - Water | 0.176464 | 0.048643 | 45 | 3.627751 | < 0.01 ** |
| Pollen & nectar - Pollen | -0.26471 | 0.059039 | 45 | -4.48369 | < 0.001 *** |
| Pollen - Water | 0.316182 | 0.053734 | 45 | 5.884198 | < 0.001 *** |
| **Species = *A. cerana,* Landscape = Forest** | | | | | |
| Empty - Pollen | 0.228269 | 0.053478 | 45 | 4.268432 | < 0.01 ** |
| Empty - Water | 0.228266 | 0.053479 | 45 | 4.268348 | < 0.01 ** |
| Nectar - Pollen & nectar | 0.336953 | 0.07574 | 45 | 4.448814 | < 0.001 *** |
| Nectar - Pollen | 0.445651 | 0.060253 | 45 | 7.396339 | < 0.001 *** |
| Nectar - Water | 0.445648 | 0.060253 | 45 | 7.396236 | < 0.001 *** |
| **Species = *A. cerana,* Landscape = Agricultural** | | | | | |
| Empty - Nectar | -0.26241 | 0.059037 | 45 | -4.44483 | < 0.001 *** |
| Nectar - Pollen & nectar | 0.248226 | 0.06031 | 45 | 4.115864 | < 0.01 ** |
| Nectar - Water | 0.354605 | 0.049274 | 45 | 7.196543 | < 0.001 *** |
| Pollen - Water | 0.16311 | 0.042107 | 45 | 3.87374 | < 0.01 ** |
| **Species = *A. cerana,* Landscape = Urban** | | | | | |
| Empty - Nectar | -0.34559 | 0.072327 | 45 | -4.77817 | < 0.001 *** |
| Empty - Pollen & nectar | 0.198527 | 0.042647 | 45 | 4.65515 | < 0.001 *** |
| Empty - Pollen | 0.169117 | 0.045984 | 45 | 3.677758 | < 0.01 ** |
| Empty - Water | 0.191173 | 0.043519 | 45 | 4.39288 | < 0.001 *** |
| Nectar - Pollen & nectar | 0.544119 | 0.049726 | 45 | 10.94226 | < 0.001 *** |
| Nectar - Pollen | 0.514709 | 0.054018 | 45 | 9.52848 | < 0.001 *** |
| Nectar - Water | 0.536765 | 0.050845 | 45 | 10.55685 | < 0.001 *** |
| **Species = *A. florea,* Landscape = Forest** | | | | | |
| Empty - Nectar | -0.22448 | 0.062122 | 45 | -3.61353 | < 0.01 ** |
| Empty - Pollen & nectar | -0.31632 | 0.065494 | 45 | -4.82972 | < 0.001 *** |
| Nectar - Water | 0.316337 | 0.049141 | 45 | 6.437313 | < 0.001 *** |
| Pollen & nectar - Pollen | 0.275514 | 0.070372 | 45 | 3.915107 | < 0.01 ** |
| Pollen & nectar - Water | 0.408175 | 0.051701 | 45 | 7.894879 | < 0.001 *** |
| Pollen - Water | 0.132661 | 0.037177 | 45 | 3.568357 | < 0.05 * |
| **Species = *A. florea,* Landscape = Agricultural** | | | | | |
| Empty - Nectar | -0.18518 | 0.058777 | 45 | -3.15059 | < 0.05 * |
| Empty - Pollen | -0.28703 | 0.062869 | 45 | -4.56558 | < 0.001 *** |
| Nectar - Water | 0.27777 | 0.04691 | 45 | 5.921293 | < 0.001 *** |
| Pollen & nectar - Pollen | -0.22222 | 0.069701 | 45 | -3.18824 | < 0.05 * |
| Pollen & nectar - Water | 0.157398 | 0.039636 | 45 | 3.971029 | < 0.01 ** |
| Pollen - Water | 0.37962 | 0.050236 | 45 | 7.55669 | < 0.001 *** |
| **Species = *A. florea,* Landscape = Urban** | | | | | |
| Empty - Pollen & nectar | 0.299062 | 0.057936 | 45 | 5.161915 | < 0.001 *** |
| Empty - Pollen | 0.214951 | 0.067429 | 45 | 3.187798 | < 0.05 * |
| Empty - Water | 0.31775 | 0.055446 | 45 | 5.73083 | < 0.001 *** |
| Nectar - Pollen & nectar | 0.261685 | 0.056596 | 45 | 4.623693 | < 0.001 *** |
| Nectar - Water | 0.280373 | 0.054165 | 45 | 5.176279 | < 0.001 *** |
| **CONTRAST: SPECIES (within each role)** | | | | | |
| **Role = Empty** | | | | | |
| *A. cerana - A. florea* (**Forest**) | 0.169693 | 0.055553 | 45 | 3.054598 | < 0.05 * |
| *A. mellifera - A. cerana* (**Urban**) | -0.1397 | 0.045058 | 45 | -3.1004 | < 0.01 ** |
| *A. mellifera - A. florea* (**Urban**) | -0.27088 | 0.053542 | 45 | -5.05929 | < 0.001 *** |
| **Role = Nectar** | | | | | |
| *A. mellifera - A. cerana* (**Forest**) | -0.30545 | 0.065162 | 45 | -4.68761 | < 0.001 *** |
| *A. mellifera - A. cerana* (**Agricultural**) | -0.15663 | 0.05923 | 45 | -2.64447 | < 0.05 * |
| *A. mellifera - A. cerana* (**Urban**) | -0.32354 | 0.056676 | 45 | -5.7086 | < 0.001 *** |
| *A. cerana - A. florea* (**Urban**) | 0.251786 | 0.062189 | 45 | 4.048734 | < 0.001 *** |
| **Role = Pollen & nectar** | | | | | |

| | | | | | |
|---|---|---|---|---|---|
| *A. cerana - A. florea* (**Forest**) | -0.2662 | 0.062333 | 45 | -4.27054 | < 0.001 *** |
| *A. mellifera - A. cerana* (**Urban**) | 0.095585 | 0.034552 | 45 | 2.766458 | < 0.05 * |
| **Role = Pollen** | | | | | |
| *A. cerana - A. florea* (**Agricultural**) | -0.17829 | 0.05861 | 45 | -3.04201 | < 0.05 * |
| *A. mellifera - A. cerana* (**Urban**) | 0.330889 | 0.047665 | 45 | 6.941969 | < 0.001 *** |
| *A. mellifera - A. florea* (**Urban**) | 0.245541 | 0.054954 | 45 | 4.468083 | < 0.001 *** |
| **Role = Water** | | | | | |
| *A. mellifera - A. cerana* (**Forest**) | 0.191225 | 0.0478 | 45 | 4.000494 | < 0.001 *** |
| *A. mellifera - A. florea* (**Forest**) | 0.22451 | 0.043998 | 45 | 5.10278 | < 0.001 *** |
| *A. mellifera - A. florea* (**Agricultural**) | 0.075813 | 0.031215 | 45 | 2.42874 | < 0.05 * |

**Table S8:** Pairwise comparisons of GRS across species for each landscape (estimated marginal means from the best-fit CLMM, ΔAICc = 0). P-values adjusted using Tukey method. SE denotes standard error, and the Z-ratio is the test statistic (ratio between estimate and standard error).

| **Contrast** | **Estimate** | **SE** | **Z-ratio** | **P-value** |
|---|---|---|---|---|
| **Landscape = FOREST:** | | | | |
| *A. mellifera - A. cerana* | 1.780 | 0.220 | 8.090 | < 0.001*** |
| *A. mellifera - A. florea* | 3.046 | 0.270 | 11.293 | < 0.001*** |
| *A. cerana - A. florea* | 1.267 | 0.275 | 4.612 | < 0.001*** |
| **Landscape = AGRICULTURAL:** | | | | |
| *A. mellifera - A. cerana* | 1.032 | 0.200 | 5.148 | < 0.001*** |
| *A. mellifera - A. florea* | 2.765 | 0.363 | 7.607 | < 0.001*** |
| *A. cerana - A. florea* | 1.733 | 0.357 | 4.857 | < 0.001*** |
| **Landscape = URBAN:** | | | | |
| *A. mellifera - A. cerana* | 0.967 | 0.205 | 4.724 | < 0.001*** |
| *A. mellifera - A. florea* | 3.325 | 0.325 | 10.220 | < 0.001*** |
| *A. cerana - A. florea* | 2.358 | 0.322 | 7.317 | < 0.001*** |
